# Variations in cell-surface ACE2 levels alter direct binding of SARS-CoV-2 Spike protein and viral infectivity: Implications for measuring Spike protein interactions with animal ACE2 orthologs

**DOI:** 10.1101/2021.10.21.465386

**Authors:** Soheila Kazemi, Alberto Domingo López-Muñoz, Jaroslav Hollý, Ling Jin, Jonathan W. Yewdell, Brian P. Dolan

## Abstract

The severe acute respiratory syndrome coronavirus 2 (SARS-CoV-2) is the causative agent of COVID-19, the most severe pandemic in a century. The virus gains access to host cells when the viral Spike protein (S-protein) binds to the host cell-surface receptor angiotensin-converting enzyme 2 (ACE2). Studies have attempted to understand SARS-CoV-2 S-protein interaction with vertebrate orthologs of ACE2 by expressing ACE2 orthologs in mammalian cells and measuring viral infection or S-protein binding. Often these cells only transiently express ACE2 proteins and levels of ACE2 at the cell surface are not quantified. Here, we describe a cell-based assay that uses stably transfected cells expressing ACE2 proteins in a bi-cistronic vector with an easy to quantify reporter protein to normalize ACE2 expression. We found that both binding of the S-protein receptor-binding domain (RBD) and infection with a SARS-CoV-2 pseudovirus is proportional to the amount of human ACE2 expressed at the cell surface, which can be inferred by quantifying the level of reporter protein, Thy1.1. We also compared different ACE2 orthologs which were expressed in stably transfected cells expressing equivalent levels of Thy1.1. When ranked for either viral infectivity or RBD binding, mouse ACE2 had a weak to undetectable affinity for S-protein while human ACE2 was the highest level detected and feline ACE2 had an intermediate phenotype. The generation of stably transfected cells whose ACE2 level can be normalized for cross-ortholog comparisons allows us to create a reusable cellular library useful for measuring emerging SARS-CoV-2 variant’s ability to potentially infect different animals.

**Importance:** SARS-CoV-2 is a zoonotic virus responsible for the worst global pandemic in a century. An understanding of how the virus can infect other vertebrate species is important for controlling viral spread and understanding the natural history of the virus. Here we describe a method to generate cells stably expressing equivalent levels of different ACE2 orthologs, the receptor for SARS-CoV-2, on the surface of a human cell line. We find that both binding of the viral Spike protein receptor binding domain (RBD) and infection of cells with a SARS-CoV-2 pseudovirus are proportional to ACE2 levels at the cell surface. Adaptation of this method will allow for the creation of a library of stable transfected cells expressing equivalent levels of different vertebrate ACE2 orthologs which can be repeatedly used for identifying vertebrate species which may be susceptible to infection with SARS-CoV-2 and its many variants.

## Introduction

The worldwide SARS-CoV-2 pandemic gave rise to an urgent need for accurate diagnosis, effective treatment, and vaccine. Rapid progress has been made in understanding the steps of virus-cell entry, understanding the nature of the virus and its viral spread both through humans and animals. It is well established that ACE2 serves as the cell-surface receptor for both SARS-CoV-1 and SARS-CoV-2, though other molecules no doubt play an important role in viral infectivity [1-5]. Due to the zoonotic nature of SARS-CoV-2 infection, there have been numerous studies attempting to understand how different vertebrate ACE2 orthologs can interact with the S-protein of SARS-CoV-2 [6-9]. This knowledge is necessary not only to understand the natural history of the virus, but also to identify species which may be more susceptible to infection. Even though ACE2 is a well conserved protein across the vertebrate clade, human polymorphisms have been documented [10], and ACE2 orthologs from different animals have unique amino acid sequences, which potentially alter the ability of a particular S-protein to bind to specific cells [11, 12]. The difference in ACE2 orthologs is thought to be a potential mechanism by which certain species appear to be protected from infection, or how other species can act as an intermediate when SARS-like coronavirus’s transition between host species [13, 14].

Since the beginning of the current global pandemic there have been multiple studies attempting to determine which ACE2 orthologs serve as receptors for S-protein. These studies include 1.) *in silico* models that predict which orthologs would be likely to bind S-protein based upon amino acid residues known to interact with S-protein [4, 9, 15-25], 2.) *in vitro* cell-based studies where viral infectivity of cells from different animals or cell-lines genetically modified to express different ACE2 orthologs are examined for interactions with S-protein (or its RBD) or infection with the virus or S-protein expressing pseudovirus [2, 6, 11, 26-30], 3.) and biochemical studies measuring the binding of S-protein with different ACE2 proteins [31, 32]. There are advantages and drawbacks to each approach. *In silico* modeling enables rapid comparison of multiple orthologs, but is limited to publicly available genetic data and predictions need to be validated experimentally. Biochemically measuring SARS-CoV-2 S-protein and ACE2 association provides an accurate measurement of the binding affinity between the two proteins, but requires the production and purification of each ACE2 ortholog and the SARS-CoV-2 S-protein or RBD and does not necessarily reflect association in the context of physiological virus-cell interactions. Cell infectivity or binding of soluble SARS-CoV-2 S-protein can provide information about which ACE2 orthologs can successfully interact with S-protein at the cell surface. However, it requires genetically modifying cells by plasmid transfection or lentiviral transduction with each ACE2 ortholog and controlling for the level of ACE2 at the cell surface is difficult.

*In vitro* infection studies with multiple different types of viruses have demonstrated that the level of viral receptor protein expressed at the cell-surface is a critical factor for viral entry into cells. For instance, Davis *et al*. reported that cells expressing the highest levels of transfected DC-SIGN(R) are more susceptible to West Nile Virus infection [33] than cells expressing lower levels of DC-SIGN(R). Likewise, Koutsoudakis *et al*. found that entry of hepatitis C virus into cells is dependent on CD81 cell-surface expression and defined a range of CD81 expression where viral infectivity is directly proportional to CD81 expression [34]. Increasing expression of Coxsackie-adenovirus-receptor (CAR) on poorly permissive cell lines increased the susceptibility of tumor cells for recombinant adenovirus infection [35]. Human immunodeficiency virus can utilize both CD4 and CCR5 as a receptor, and the relative expression of each receptor has been shown to influence infectivity for different HIV isolates [36]. Therefore, when measuring SARS-CoV-2 infectivity *in vitro*, it is critical to consider the amount of ACE2 at the cell surface.

Here we describe the stable transfection of HRT-18G cells, a human rectal cancer cell line capable of supporting beta-coronavirus replication [37, 38], with plasmid vectors containing a bi-cistronic mRNA encoding different ACE2 orthologs and a cell-surface reporter protein, mouse Thy1.1. We demonstrate that Thy1.1 is directly proportional to the amount of human ACE2 on the cell surface and increasing ACE2 expression leads to both increased binding of a fluorescent RBD and increased SARS-CoV-2 pseudoviral infectivity. We also generated stable cell lines expressing either mouse or domestic cat ACE2 with equivalent Thy1.1 expression to the human ACE2 cells and found that RBD binding, and infectivity is absent upon expression of mouse ACE2, while feline ACE2 has an intermediate phenotype when compared to human. Our system can be easily expanded to include more ACE2 orthologs to provide further insight as to how RBD/ACE2 interactions impact the susceptibility or resilience of species against SARS-CoV-2 infection and transmission.

## Results

### ACE2 binding to SARS-CoV-2 Spike protein is necessary for viral infectivity

SARS-CoV-2 gains access to cells when its S-protein binds to the cell-surface receptor ACE2 [1-4, 39, 40]. To confirm this in our chosen cell system we generated a cell line stably expressing human ACE2 (hACE2) to examine the S-protein binding and SARS-CoV-2 pseudoviral infectivity. We generated DNA plasmids encoding a bi-cistronic mRNA containing human ACE2 cDNA followed by an internal ribosome entry site (IRES) and finally the mouse cell-surface protein Thy1.1 (Figure 1A). Because Thy1.1 and ACE2 are translated by the same mRNA, we used the relative expression of Thy1.1 as detected by widely available monoclonal antibodies to infer ACE2 expression, thus avoiding uncertainties regarding the interaction of assorted orthologs of ACE2 with available antibodies able to detect native ACE2 on the cell surface. Following this rationale, we transfected HRT-18G cells, a human rectal cancer line capable of supporting betacoronovirus replication, with the plasmid encoding hACE2 and Thy1.1 under conditions to support stable transfection. Following transfection, cells were magnetically sorted utilizing antibodies specific for Thy1.1. Three rounds of sorting were needed to generate a stable cell line, termed HRT-18G/hACE2 with >95% of cells expressing Thy1.1 (Figure 1B). In comparison to parent cells, stably transfected cells showed significant increase in Thy1.1 and ACE2 antibody staining (Figure 1B & 1C). To determine the capability of hACE2 expressed on HRT-18G cells to bind to the RBD of SARS-CoV-2 S-protein, we incubated the cells with Alexa Fluor-647 labeled RBD protein purchased from a commercial vendor and analyzed them by flow cytometry (Figure 1D). In contrast to parent cells, the RBD interacted with the hACE2 in a concentration dependent manner (Figure 1E). We also assessed the capability of hACE2 to allow the viral infectivity compared to parent cells. Both HRT-18G and HRT-18G/hACE2 cells were exposed to a (Vesicular Stomatitis Virus) VSV-derived SARS-CoV-2 pseudovirus expressing S-protein and GFP at an MOI of 10 for short, defined periods of time, washed to remove excess virus, and cultured for 16 hours. The number of infected cells was then determined by measuring GFP expression by flow cytometry. As shown in Figure 1F, a measurable percentage of infected HRT-18G/hACE2 cells was detected with as little as 5 minutes exposure to the virus inoculum, and the number of infected cells steadily increased as exposure time was lengthened. Finally, infected cells were visualized using fluorescence microscopy. HRT-18G and HRT-18G/hACE2 cells were infected with pseudovirus at a MOI of approximately 0.5. Twenty-four hours post-infection, cells were examined by fluorescence microscopy for the presence of GFP positive cells. In contrast to the parental HRT-18G cells, which showed no evidence of GFP expression, HRT-18G/hACE2 cells were readily infected with SARS-CoV-2 S-protein expressing pseudovirus (Figure 1G).

Taken together these findings show that HRT-18G cells can be stably transfected and sorted based upon Thy1.1 expression to generate a cell line expressing human ACE2 which can bind SARS-CoV-2 RBD protein and support pseudoviral infection.

**Figure 1.**
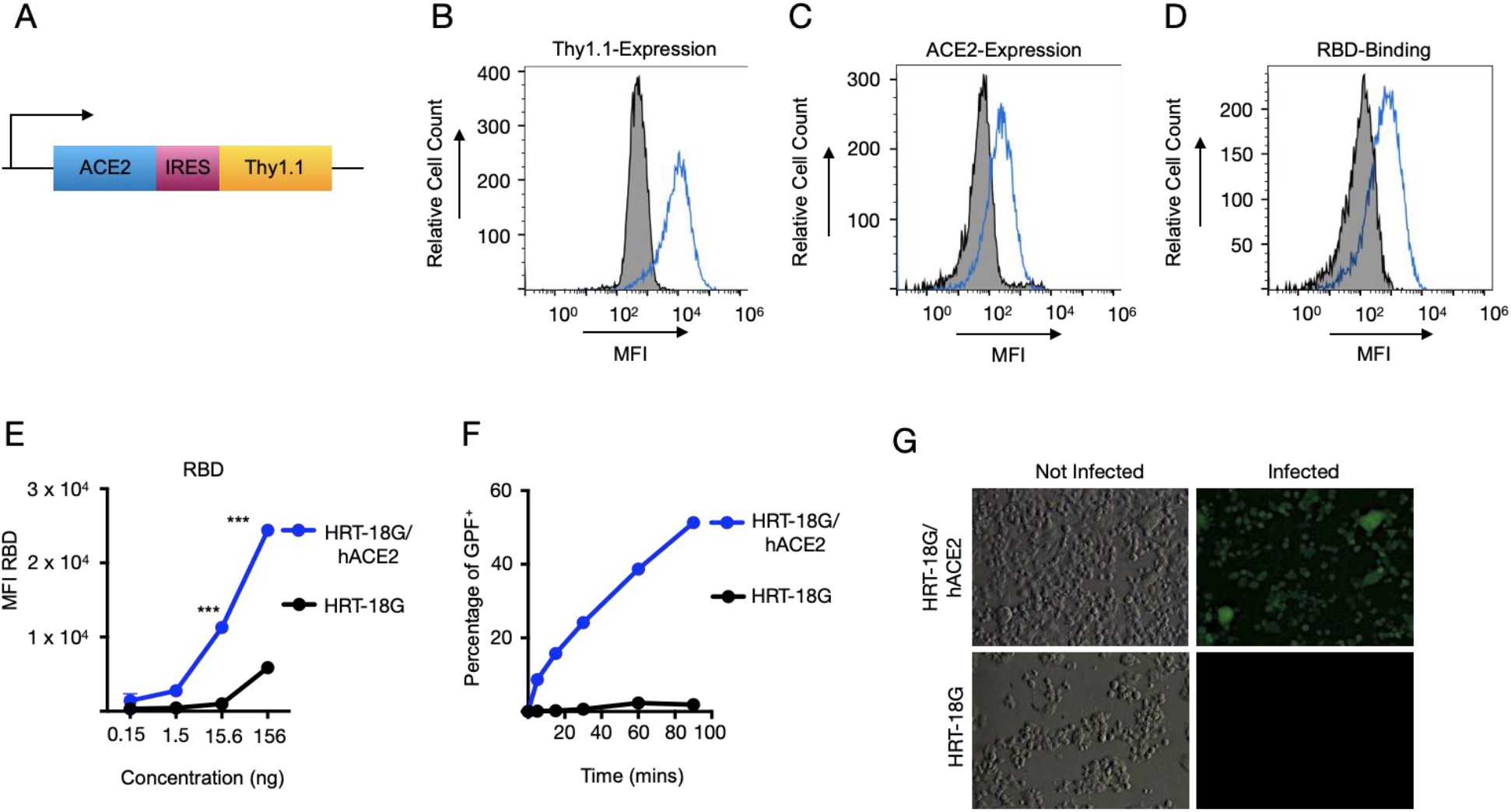
Recombinant SARS-CoV-2 RBD protein binds to human ACE2 and facilitates the pseudoviral infectivity. *A*. Schematic of the bi-cistronic mRNA for production of human ACE2 and the reporter cell surface protein, mouse Thy1.1. *B*. HRT-18G cells were stably transfected with plasmid containing cDNA of hACE2 and magnetically sorted for Thy1.1 expression. HRT18G/hACE2 (blue trace) and the parental HRT-18G (shaded histogram) cells were stained with fluorescent anti-Thy1.1 antibody and expression was analyzed by flow cytometry. *C*. HRT-18G cells stably expressing ACE2 (blue trace) and parent cells (shaded histogram) were incubated with fluorescent anti-ACE2 antibody and analyzed by flow cytometer. *D* HRT-18G parental cells (shaded histogram) and HRT-18G/hACE2 cells (blue trace) were incubated with Alexa Fluor 647-labeled SARS-CoV-2 S-protein RBD and analyzed for protein binding by flow cytometry. *E*. Same as in *D* except cells were incubated with a range of concentrations of Alexa Fluor 647-labeled SARS-CoV-2 S-protein RBD and analyzed by flow cytometer. *F*. HRT-18G/hACE2 and HRT-18G cells were infected with GFP-expressing SARS-CoV-2 pseudovirus at an MOI of 10 for the indicated times, washed to remove excess virus, and incubated at 37 °C overnight. The infectivity was analyzed by a flow cytometer 16 hours later and the percentage of GFP positive cells reported. *G*. HRT-18G/hACE2 and parent cells were infected with a GFP-expressing SARS-CoV-2 pseudovirus at an MOI of 0.5 and cultured for 24 hours prior to analysis by fluorescence microscopy. All data presented is representative of three independent experiments. Statistical significance is indicated as: *** *P < 0*.*0001*.

### Cell surface ACE2 levels govern RBD binding and viral infectivity

We hypothesized that increased expression of Thy1.1 would correlate with increased ACE2 expression leading to increased RBD binding and viral infectivity. We tested this hypothesis in two different ways. First, we transiently transfected HRT-18G cells with hACE2-IRES-Thy1.1 DNA and 48 hours later, we stained the cells for both Thy1.1 and hACE2. As shown by flow cytometry analysis (Figure 2A), Thy1.1 staining strongly correlates with ACE2 levels detected by antibody staining. As a control, we show individual Thy1.1 and ACE2 staining which result in no overlap in signal (Figure 2A). To determine the interaction of transiently expressed hACE2 with RBD of S-protein, we stained both HRT-18G and HRT-18G/hACE2 cells with Thy1.1 and fluorescently labeled SARS-CoV-2 RBD protein. Consistent with data shown in Figure 1D and 1E, a strong positive correlation was seen between Thy1.1 expression and RBD binding (Figure 2B), while staining with either Thy1.1 or fluorescent RBD protein individually demonstrated no overlap in signal.

Next, we fluorescently sorted HRT-18G/hACE2 cells based upon Thy1.1 staining to isolate the cells with the highest expression of Thy1.1, designated HRT-18G/hACE2++. As expected, Thy1.1 expression was significantly higher in these cells compared to both parent cells and HRT-18G/hACE2 cells (Figure 2C). To determine if the Thy1.1 expression is relative to the expression of ACE2 we stained HRT-18G/hACE2 and HRT-18G/hACE2^++^ cell lines with ACE2 antibodies. Consistent with the Thy1.1 expression level, HRT-18G/hACE2++ cells showed an increase in ACE2 expression compared to parent cells and HRT-18G/hACE2 (Figure 2D). HRT-18G, HRT-18G/hACE2, and HRT-18G/hACE2++ cells were then labeled with a specific concentration of fluorescent RBD and analyzed by flow cytometry. HRT-18G/hACE2++ cells showed a distinct shift in fluorescent intensity compared to parental cells and significant increase compared to HRT-18G/hACE2 cells (Figure 2E). Finally, to confirm the importance of differential ACE2 expression on viral infectivity, we conducted a time-course infection with the VSV-eGFP-deltaG-SARS-CoV-2 S-protein pseudovirus. We hypothesized that cells with more ACE2 expression at the cell surface would be more infectible when exposed to virus for brief periods of time. Consistent with the result of S-protein binding to ACE2, the percent of infected, GFP-positive cells was greater in cells expressing high levels of ACE2 (Figure 2F).

Together these data indicate that measuring Thy1.1 in our system is a good proxy for total ACE2 levels and alterations to the level of ACE2 at the cell surface can impact S-protein binding and viral infectivity.

**Figure 2.**
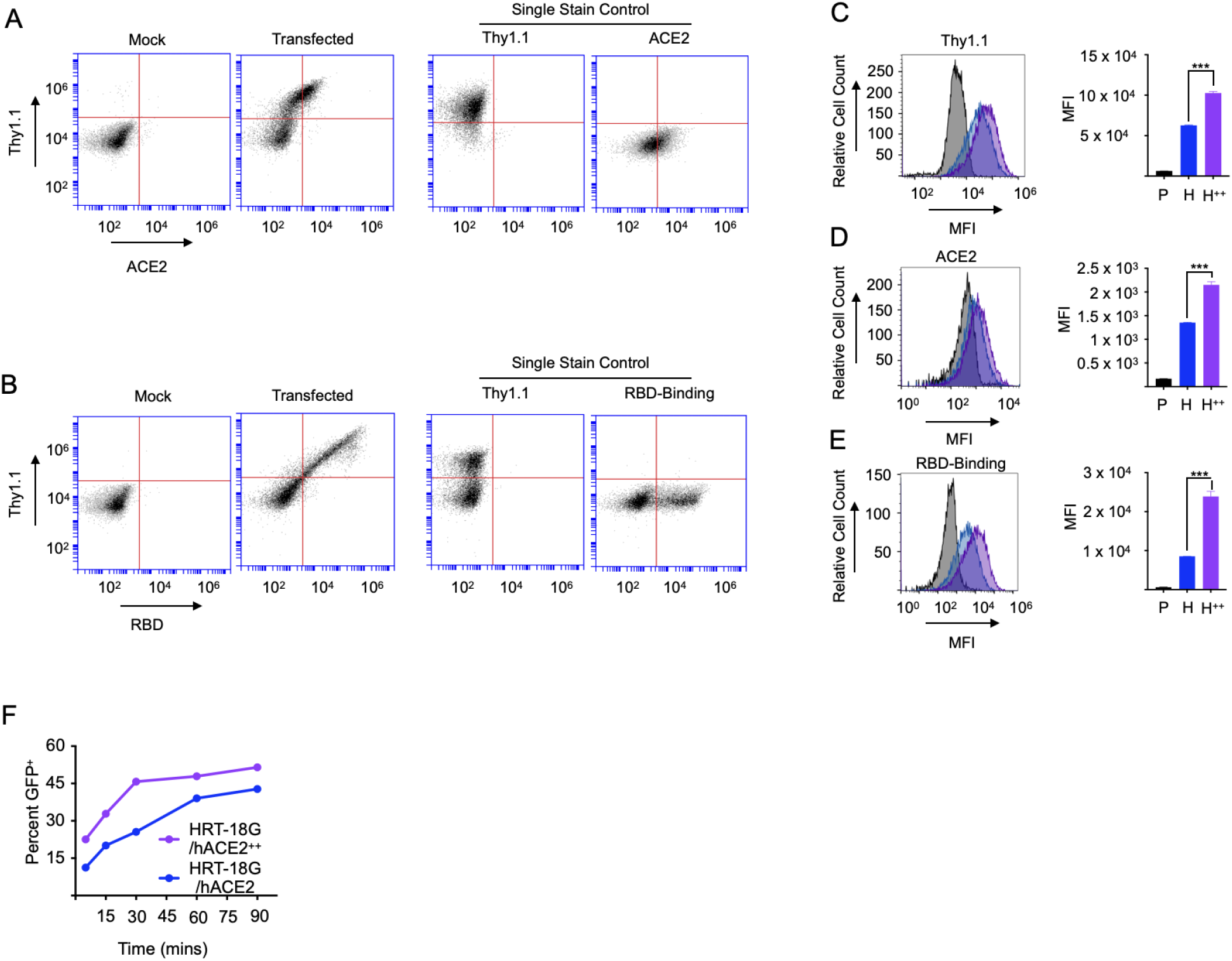
Increased ACE2 expression potentiates SARS-CoV-2 RBD binding and pseudoviral infectivity. *A, B*. HRT-18G cells were transiently transfected with cDNA encoding hACE2 and Thy1.1 in a bi-cistronic cassette and two days later stained with *(A)* fluorescently labeled anti-Thy1.1 and ACE2 antibodies and analyzed by flow cytometry. Single antibody labeling controls are also depicted. *B*. Cells were stained with fluorescently labeled anti-Thy1.1 antibody and Alexa Fluor 647-labeled SARS-CoV-2 S-protein RBD analyzed by flow cytometry. Histograms of individual protein-labeled cells are also depicted. *C*. HRT-18G/hACE2 (blue trace) were fluorescently sorted based on Thy1.1 expression to generate the cell line HRT-18G/hACE2++ (purple trace). HRT-18G/hACE2 and HRT-18G/hACE2++, and parental HRT-18G (shaded histogram) cells were then incubated with *(C)* anti-Thy1.1, *(D)* anti-ACE2 antibodies, or *(E)* Alexa Fluor 647-labeled SARS-CoV-2 S-protein RBD and analyzed by flow cytometry. Representative histograms are shown as well as quantitative MFI measurements from three technical repeats. *F*. HRT-18G/hACE2 and HRT-18G/hACE2++ cells were infected with GFP-expressing SARS-CoV-2 pseudovirus for indicated times, washed to remove excess virus, and incubated in complete media for 16 hours and analyzed by flow cytometry for GFP expression. All data presented is representative of three independent experiments. Statistical significance is indicated as: *** *P < 0*.*0001*.

### ACE2 orthologs show different degrees of SARS-CoV-2 Spike protein binding

Given the zoonotic nature of SARS-CoV-2 [8, 41], it is necessary to reliably predict the susceptibility of both wild and domestic animals to SARS-CoV-2 infection, which is controlled in part by the affinity of S-protein for ACE2. Our data indicate that the level of ACE2 at the cell surface can influence both binding of the RBD of the SARS-CoV-2 S-protein as well as viral infectivity. Therefore, studies comparing different vertebrate ACE2 orthologs need to account for the relative levels of ACE2 expressed at the cell surface.

We created two additional HRT-18G cell lines expressing either domestic feline or mouse ACE2 orthologs using the Thy1.1 bi-cistronic expression system. When compared to HRT-18G/hACE2 cells, both HRT-18G/fACE2 (feline ACE2) and HRT-18G/mACE2 (murine) expressed equivalent levels of Thy1.1 (Figure 3A) and therefore expressed equivalent levels of each ACE2 ortholog. Cells were stained with antibodies for human ACE2 (as used in Figures 1 and 2) and only HRT-18G/hACE2 cells stained positive for ACE2 (Figure 3B), indicating that this particular monoclonal antibody does not recognize either feline or mouse ACE2. We then tested for SARS-CoV-2 RBD interactions with each individual cell line by incubating fluorescently labeled RBD protein with cells and measuring fluorescent protein binding by flow cytometry. Even though levels of Thy1.1 were equivalent in all three cell lines (Figure 3A), the S-protein RBD binding varied depending on the ACE2 ortholog expressed (Figure 3C). No interaction between mouse ACE2 and S-protein was detected, and the RBD-binding pattern was the same as in parental HRT-18G cells (Figure 3C & 3D). Human ACE2 had the highest affinity against the S-protein RBD, while the feline ACE2 fell between mouse and human (Figure 3C & 3D). To further confirm the hierarchy of the S-protein RBD affinity against ACE2, we next infected the same set of cell lines with SARS-CoV-2 pseduovirions and measured the number of infected cells expressing GFP. Consistent with the S-protein RBD binding with ACE2 in figure 3D and compared to parent cells, HRT-18G/hACE2 had the most infected cells and HRT-18G/mACE2 the least infected cells, while HRT-18G/fACE2 cells ranked in between human and mouse (Figure 3E).

**Figure 3.**
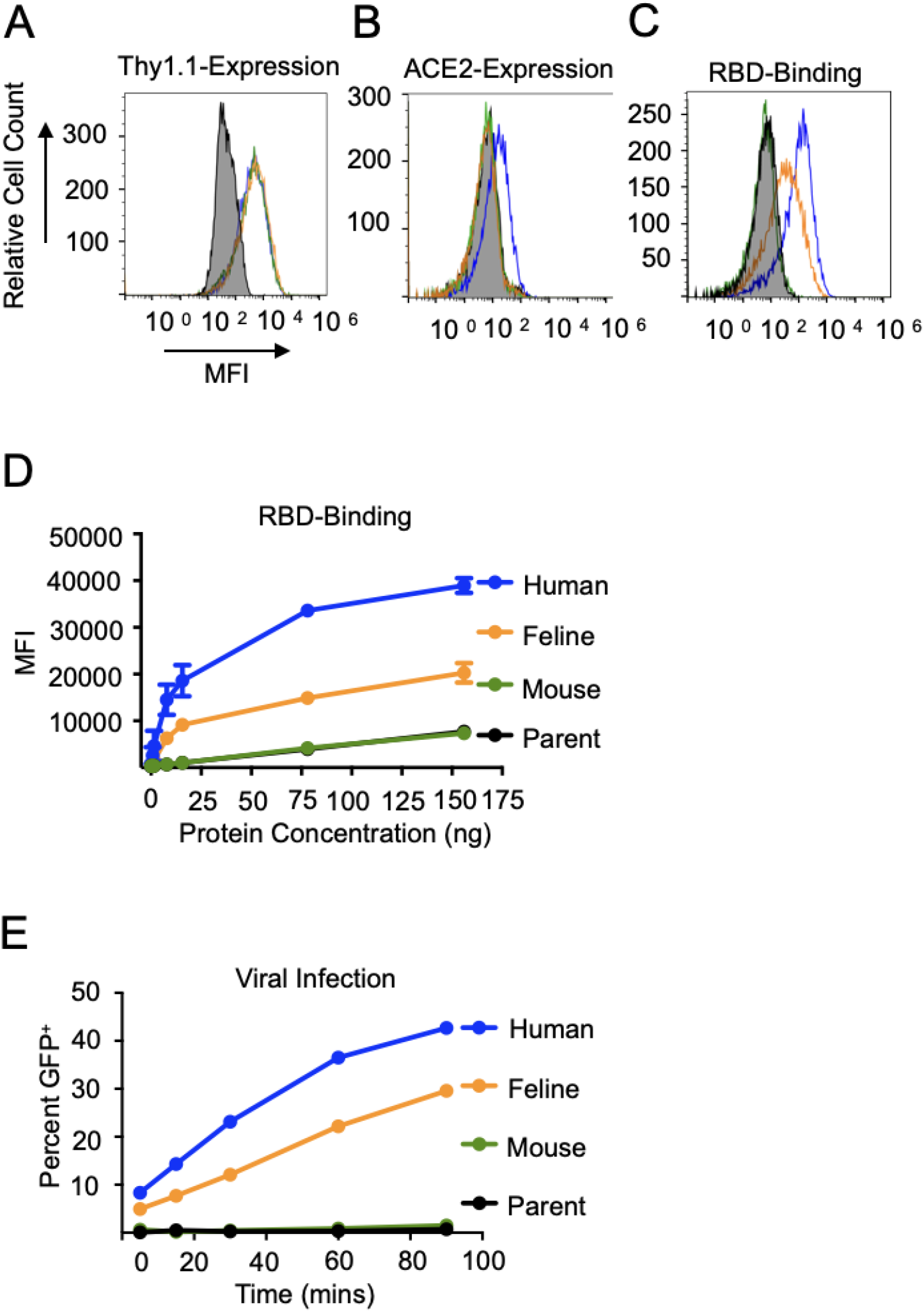
Equivalent expression of ACE2 orthologs and varied spike protein binding. *A*. HRT-18G cells were stably transfected with IRES-Thy1.1 plasmids containing cDNAs of human (blue), feline (orange), and mouse (green) ACE2 and magnetically sorted until an equivalent level of reporter protein Thy1.1 was expressed on all three cell lines. *B*. To confirm the specificity of the ACE2 antibody for human ACE2, HRT-18G/hACE2, HRT-18G/fACE2, HRT-18G/mACE2 and HRT-18G (shaded histogram) cells were simultaneously incubated with fluorescently-labeled ACE2 antibody and analyzed by flow cytometry. *C*. The four cell lines incubated with 7.8 ng of Alexa Fluor 647-labeled SARS-CoV-2 S-protein RBD and analyzed for SARS-CoV-2 S-protein RBD/ACE2 binding affinity by flow cytometry. *D*. Same as in *C* except cells were incubated with different concentrations of Alexa Fluor 647-labeled RBD. The MFI of the population is reported on the y-axis. *E*. HRT-18G/hACE2, HRT-18G/fACE2, HRT-18G/mACE2 and HRT-18G cells were infected with GFP-expressing SARS-CoV-2 pseudovirus for specified times, washed to remove excess virus, and incubated for 16 hours, and GFP expressing cells quantified by flow cytometry. All data presented is representative of three independent experiments.

## Discussion

The use of stably transfected cells expressing equivalent levels of assorted ACE2 orthologs provides a number of advantages. Of particular importance is the expression of a defined amount of ACE2 which can be achieved through the combination of stable transfection along with expression of a reliable reporter protein. As we show in Figure 2, levels of easily-measurable Thy1.1 are proportional to ACE2 levels, which in turn, correlates with both SARS-CoV-2 S-protein RBD binding and S-protein-mediated viral infectivity. In systems with transient expression of ACE2 genes [6, 27, 30, 42], levels of ACE2 vary not only within a given population of cells, but also between individual experiments. For example, as shown in Figure 2, transient expression of hACE2 reveals a broad level of ACE2 expression which directly impacts S-protein RBD binding. Selecting cells based upon defined levels of Thy1.1 expression ensures consistent levels of ACE2. This is important when comparing the interaction of S-protein with different ACE2 orthologs or homologs.

Several studies to date have expressed different ACE2 orthologs on cells to measure the potential for SARS-CoV-2 infectivity. The results have shown some consistent patterns, such as the inability of *Mus musculus* ACE2 to interact with SARS-CoV-2 [2, 24, 26, 32], (with the exception of mouse cells from the cornea [43]) or the relatively high infectivity afforded by pangolin ACE2 [6, 26, 28, 29]. Results from other orthologs are more inconsistent. Conceicao *et al*. noted discrepancies in studies examining a variety of bat ACE2 orthologs expressed by non-permissive cells [28]. In some reports, horseshoe bat ACE2 expression leads to robust viral infectivity or S-protein binding [2, 4, 26] whereas other reports suggest a more modest or very weak binding and infectivity [11, 27-31]. Similar discrepancies appeared when examining palm civet ACE2, with some reports indicating poor infectivity and/or S-protein binding [11, 28-32] in contrast with other reports [2, 26]. Variations in cell surface ACE2 levels between the cells used to make these comparisons likely explain some of these discrepancies. The Thy-1.1 reporter system we describe may be useful in resolving these findings.

Another advantage of our system is allowing researchers to infer cell-surface levels of ACE2 without the need for species-specific reagents. As shown in Figure 3, a monoclonal antibody that recognizes human ACE2 fails to stain cells expressing either feline or mouse ACE2, demonstrating that commercially available reagents may not be useful for measuring the cell-surface levels of different ACE2 orthologs. Others have used polyclonal antibodies to detect different ACE2 orthologs [6], however, it is not possible to determine precise levels of ACE2 with polyclonal sera as the mixture of different antibodies with different affinities for different ACE2 orthologs confounds the ability to quantify expression. These reagents are useful in determining if ACE2 is expressed but cannot reliably determine relative amounts of different ACE2 orthologs. The use of a reporter in a bi-cistronic vector that can easily be measured, and used for fluorescence-based cell sorting, allows us to compare SARS-CoV-2 S-protein interactions and viral infectivity without the need to develop and characterize specific reagents for each ACE2 ortholog to be studied.

Stably transfected cells can also be cultured indefinitely, which negates the need to repeatedly transfect cells for each new experiment. The necessity of these renewable cells is especially important when considering newly emerging SARS-CoV-2 variants. Many of the variants of concern have altered binding of the viral S-protein to ACE2, which is of great concern to public health [44-47]. It is still unknown if the newly emerged variants are more infectious to other vertebrate animals, or if there is a shift of a species predicted susceptibility to a newly emerged variant. With the presence of stably transfected cells expressing a defined amount of ACE2 that simply need to be thawed from cryogenic storage, researcher will be able to rapidly determine if particular variants are more or less infectious to particular animals. It should be noted that measuring ACE2 interactions with SARS-CoV-2 may not be a perfect method for predicting a species susceptibility to infection, progression to disease, and even viral transmission. For instance, both *in silico* and *in vitro* studies have suggested that ferrets and mink ACE2 interacts poorly with S-protein [6, 9, 48], however these animals are readily infectible with SARS-CoV-2, often dramatically so [49-52]. As the field progresses, it will behoove us to think not only of ACE2 but other physiological differences between species which may impact SARS-CoV-2 susceptibility.

## Materials and Methods

### Cell lines, antibodies, and reagents

HRT-18G and BHK-21 cells were obtained from American Type Culture Collection (ATCC, Manassas, VA) and authenticated by the provider. Cell cultures were maintained in Dulbecco’s Modified Eagle’s Medium (DMEM, Gibco, MA, USA) supplemented with 1% GlutaMax (Gibco), 7.5% fetal bovine serum (Atlanta Biological Inc., GA, USA), at 37 °C in a humidified incubator containing 6% CO2. Cell lines were checked regularly for any mycoplasma contamination and were tested negative for the presence of mycoplasma contamination using Universal Mycoplasma Detection Kit (ATCC^®^ 30-1012K™). Cell lines and their derivative cell lines were expanded and frozen in a liquid nitrogen tank in cryogenic storage at low passage number. All Thy1.1 antibodies (unlabeled, and FITC conjugated) were from eBiosciences and APC-coupled anti-ACE2 antibody was from Abeomics. All antibodies were used according to manufacturer’s protocols. Alexa Fluor 647-protein labeling kit was from Invitrogen.

### Plasmid generation for ACE2 expression

All primers were obtained from Integrated DNA Technologies and PCR products were amplified using a Veriti thermocycler (Applied Biosystem). For the generation of ACE2 expressing BHK-21 cells, human ACE2 cDNA (kindly provided by Sonja Best, NIH/NIAID) was used as a PCR template. The open reading frame of hACE2 was amplified in PCR using hACE2_F: 5’-TGGCCTGACAGGCCCTAAAAGGAGGTCTGAACATCATC-3’ and hACE2_R: 5’-TGGCCTCTGAGGCCACCATGTCAAGCTCTTCCTGGC-3’ primers and cloned to pSBbi-BH vector using *SfiI* restriction site. The pSBbi-BH plasmid was a kind gift from Eric Kowarz (Addgene #60515). ACE2 orthologs were cloned into the plasmid pCAGGS-IRES-Thy1.1, as described previously [53]. ACE2 cDNA sequences encoding human, feline, or murine ACE2 were ordered as gene blocks (IDT) using the NCBI reference sequences NM_001371415.1, NM_001039456.1, and NM_001130513.1 respectively. Gene blocks were appended with either an NheI or ClaI site at the N-terminus and AgeI site on the C-terminus. Gene blocks fragments and pCAGGS-IRES-Thy1.1 were digested with appropriate restriction enzymes, gel purified, ligated together using Mighty Mix ligation kit (Takara), cloned into competent DH5α *E. coli* and selected for ampicillin resistance. ACE2 sequences in positive clones were confirmed by Sanger DNA sequencing at the Oregon State University Center for Quantitative Life Sciences (CQLS). Plasmids were purified using Qiagen HiSpeed Midi Plasmid Purification kit according to the manufacturer’s instructions.

### Cell transfection and stable cell line generation

For the generation of BHK-21 cells stably expressing human ACE2, the pCMV(CAT)T7-SB100 plasmid (encoding transposase required to generate stable transgene-expressing cell lines) was co-transfected with pSBbi-BH hACE2 using TransIT-LT1 Transfection Reagent (Mirus Bio). pCMV(CAT)T7-SB100 was a kind gift from Zsuzsanna Izsvak (Addgene #34879). The cells were selected with 250 mg/ml hygromycin B Gold (InvivoGen) for 2 weeks. Surface expression of hACE2 was confirmed by flow cytometry using anti-hACE2 antibody. For HRT-18G cell lines, 2.2 μg of pCAGGS-IRES-Thy1.1 vectors encoding ACE2 orthologs were linearized with PvuI, subsequently purified, and 10^6 cells were transfected into cells using Amaxa 96-well Nucleofector (Lonza, Basel, Switzerland) SF kit on program DS-138. Transfected cells were kept at 37 °C until optimal confluency was obtained. Cells were then prepared for magnetic bead sorting using the MidiMACS™ Separator according to manufacturer’s instructions. Briefly, cells were harvested and washed in specific sorting buffer, incubated with anti-Thy1.1 antibody for 30 min at 4°C. Cells were washed again and incubated with anti-mouse IgG microbeads (Miltenyi Biotech, Bergisch Gladbach, Germany) for 15 min at 4 °C. Washed cells were passed through the LS column, eluted cells were mixed with fresh media and kept at 37 °C. The process was repeated until a population of more than 90% Thy1.1 expression was acquired.

### RBD labeling

Recombinant SARS-CoV-2 Spike RBD (derived from strain Wuhan-Hu-1) protein was purchased from ABclonal Technology and labeled with Alexa Fluor 674 reactive fluorescence dye according to manufacturer’s protocols (Invitrogen). HRT-18G cells and their derivatives were labeled by incubating the cells with a specific concentration of the labeled RBD for 30 min at 4 °C. Cells were then washed with Hank’s balanced salt solution (HBSS) supplemented with 0.1% BSA and were resuspended in 0.1% BSA/HBSS and analyzed with Accuri C6 benchtop flow cytometer (BD Biosciences, San Jose, CA, USA).

### Flow cytometry

HRT-18G cells stably or transiently expressing ACE2 orthologs were harvested, washed with HBSS supplemented with 0.1% BSA. Cells were then labeled with FITC coupled anti-Thy1.1, APC coupled anti-ACE2 or isotype control and incubated for 30 min at 4 °C. Cells were resuspended in 0.1% BSA/HBSS and analyzed with Accuri C6 benchtop flow cytometer (BD Biosciences, San Jose, CA, USA). For GFP analysis of virally infected cells, cells were harvested, washed in PBS, fixed in 2% paraformaldehyde for 15 minutes at room temperature, washed in excess PBS, and analyzed for GFP expression.

### Virus generation, propagation, and infection

The generation of VSV-eGFP-deltaG_SARS-CoV-2 S-protein (hereafter SARS-CoV-2 pseudovirus) was as follows. The SARS-CoV-2 Spike full length sequence was amplified from vector # NR-52310 (BEI) with primers TGTTTCCTTGACACTATGTTCGTGTTTCTGGTGCTG and CACAAGTTGATTTGGTCAGGTGTAGTGCAGTTTCAC. The backbone vector pVSV-eGFP-dG (Addgene # 31842) was linearized by PCR with primers AGTGTCAAGGAAACAGATCGATCTC and CCAAATCAACTTGTGATATCATGC. The SARS-CoV-2 Spike sequence was cloned into the linearized pVSV-eGFP-dG by the In-Fusion cloning system (Takara Bio Inc.). The resulting pVSV-eGFP-deltaG_SARS-CoV-2 Spike vector was verified by sanger sequencing. VSV-eGFP-deltaG_SARS-CoV-2 Spike was recovered as previously described [54]. Briefly, BHK-21 and 293FT cells were plated together in 6-well plates and infected with recombinant T7-expressing vaccinia virus (vTF7-3). Virus inoculum was removed after 1 h and cells were transfected with a mixture of plasmids using TransIT-LT1 (Mirus Bio # MIR 2300), including pVSV-eGFP-deltaG_SARS-CoV-2 Spike (5 µg), N (Addgene # 64087), P (Addgene # 64088), and L (Addgene # 64085) recovery support plasmids (3:5:1 µg). At 48 h after transfection, the supernatant was collected, filtered through a 0.22 µm filter to remove vTF7-3, and used to infect fresh BHK-21/hACE2-BFP in 6-well plates. The recovered virus was identified by fluorescence microscopy screening, passaged one more time in BHK-21/hACE2-BFP and then, amplified by infecting BHK-21/hACE2-BFP in 75-cm^2^ flaks. After 72 h post infection, the supernatant was collected, cell-debris clarified, aliquoted and frozen at −80ºC. The presence of SARS-CoV-2 Spike was confirmed by western blot with anti-SARS-CoV-2 S1 (Sino Biological # 40150-D004) and anti-SARS-CoV-2 S2 (Thermo Fisher # PA5-112048). The inability of the rescued virus to infect BHK-21 cells in the absence of hACE2 was assessed by fluorescence microscopy and flow cytometry. The virus was propagated by infecting BHK-21/hACE2-BFP cells grown to confluency in a T-125 cm2 flask with a low dose of virus (equivalent to an approximate MOI of 0.1) diluted in media to 15 ml in media containing 2% FCS for 48 hours. The flask was then subject to three rounds of freeze-thaw to lyse cells and the media centrifuged at 300 RCF for 15 minutes to pellet insoluble debris. Virus-containing supernatant was aliquoted and stored at −80 C. Virus was tittered on BHK-21/hACE2-BFP cells and counting visible plaques 72 hours post infection. For infection studies, HRT-18G cells and their derivatives were plated in 24 well plates, 24 hours prior to infection to achieve confluency. Media was removed and virus containing supernatant was added to the cells. For kinetic infections, virus was added at an (Multiplicity of Infection) MOI of ∼10 and cells were incubated for the indicated time. The virus-containing media was removed, and cells were washed with complete media, and then incubated for 16 hours in complete media.

### Analysis of GFP-VSV infection by high-content imaging system

BHK-21/hACE2-BFP cells were infected with VSV-eGFP-deltaG_SARS-CoV-2 S protein pseudovirus at an MOI of 0.5 for 2 hours with occasional agitation. Cells were then incubated with virus-free complete media and incubated at 37 °C for 24 hours. The next day, cells were analyzed by fluorescence microscopy (Leica DMIL LED), using Qcapture Pro 6.0 inverted scope camera to capture cellular images.

### Statistical analysis

All statistical analyses were performed using GraphPad Prism Software version 9.0 (GraphPad Software Inc., San Diego, CA). Multiple groups comparisons were performed through one-way analysis of variance (ANOVA), followed by Tukey’s or Dunnett’s multiple comparison post hoc test. Values of p < 0.05 were considered statistically significant.

